# Single-particle light scattering reveals structural heterogeneity of biomolecular condensates

**DOI:** 10.64898/2026.04.10.717862

**Authors:** Berenice García Rodríguez, Katarzyna Makasewicz, Giulio Tesei, Fredrik Eklund, Edvin Johansson, Paolo Arosio, Giovanni Volpe, Daniel Sundås Midtvedt

## Abstract

Biomolecular condensates can be heterogeneous in composition, morphology, and material properties, but resolving this heterogeneity at the level of individual submicrometer condensates remains challenging. Here, we combine off-axis holographic imaging with quantitative light-scattering analysis and particle tracking to simultaneously measure the size, optical properties, interfacial structure, and hydrodynamic mobility of hundreds of individual condensates per minute. Applied to condensates formed by the N-terminal domain of Ddx4 (Ddx4N1), the method reveals two populations with distinct scattering signatures: compact particles with well-defined interfaces and structurally heterogeneous, non-compact assemblies. Their relative abundance varies systematically with ionic strength. Synthetic polymer controls show that pronounced structural heterogeneity is not a generic consequence of phase separation and suggest a role for interaction heterogeneity in promoting competing mesoscale structures. Mass–size scaling and Brownian aggregation simulations further show that aggregation with incomplete fusion of smaller protein-rich units provides a physical model for the non-compact population. Simultaneous optical and diffusion measurements further show that the two populations have distinct hydrodynamic properties. These results demonstrate that chemically homogeneous condensate-forming systems can populate distinct mesoscale structures that are obscured by ensemble-averaged measurements.

## Introduction

Biomolecular condensates are membraneless assemblies that arise through weak, multivalent interactions among proteins and nucleic acids [1–3]. Their physical properties, including size, internal composition, interfacial structure, and morphology influence their material state, biochemical activity, and responsiveness to environmental cues. Furthermore, these properties are dynamic, with important functional implications [4]. For example, initially liquid-like condensates may coarsen, for instance through percolation [5] or amyloid fibril formation [6], into more solid-like structures, a process linked to a variety of diseases [7]. Furthermore, the composition and structure of condensates can be modulated by biochemical reactions [8] and post-translational modifications [9]. Quantitatively resolving the physical mesoscale properties of condensates is therefore essential for understanding their formation and function.

There has been significant progress in recent years in the development of methods for condensate characterization [10]. Existing approaches have provided important insights into condensate size, dynamics, and material properties, particularly for large, micrometer-scale condensates formed at high protein concentrations. However, most existing single-condensate studies probe only a small number of large condensates, yielding limited statistics. In contrast, condensates formed under biologically relevant conditions are frequently submicrometer in size and exhibit weak refractive index contrast, which complicates their quantitative characterization by conventional microscopy approaches.

Recent developments in optical single-nanoparticle characterization have expanded the ability to probe individual structures at the submicrometer scale [11–16]. However, most available methods estimate particle size from diffusion, which requires assumptions about the relationship between mobility and physical size that do not hold for soft, permeable condensates. At even smaller scales, protein assemblies can be characterized on the nanometer scale using ensemble-based approaches, such as dynamic light scattering (DLS) [17] or small-angle x-ray scattering (SAXS) [18], but such techniques will inherently obscure sample heterogeneity. Consequently, a high-throughput, multiparametric technique for characterizing submicrometer condensates and resolving their structural heterogeneity is still lacking.

Here, we introduce an interferometric single-particle scattering approach for high-throughput characterization of submicrometer biomolecular condensates. By combining Fourier-domain analysis of scattering patterns with three-dimensional particle tracking, the method independently probes particle size, optical properties, interfacial structure, and hydrodynamic mobility. We apply this approach to Ddx4N1 condensates and synthetic polymer model systems. The measurements reveal substantial particle-to-particle heterogeneity that is not captured by size alone, including distinct scattering and hydrodynamic signatures whose prevalence varies across phase-separation conditions. By combining these observables, we distinguish compact condensates from structurally heterogeneous assemblies and investigate physical models capable of accounting for their mesoscale architecture.

## Results

### Single-particle light scattering quantifies size and refractive index of submicrometer particles

The off-axis holographic imaging platform used in this study is outlined in Figure 1. We employed an off-axis holographic microscope (Fig. 1**a**) to record submicrometer particles flowing through a microfluidic channel. The three-dimensional tracking enabled by holographic imaging (Fig. 1**b**) allows reconstruction of the complex scattered field for each particle as it passes the microfluidic channel. For each trajectory, we obtain a stack of numerically refocused images containing the real and imaginary components of the scattered field. Fourier transformation of this complex field, followed by calculation of its magnitude, yields the scattering amplitude in the back focal plane of the objective. In this representation, the signal directly reports the distribution of the scattered light across the entrance pupil and thus provides a quantitative description of its angular scattering profile (Fig. 1**c**; see Methods and Supporting Information). Because each particle is observed over typically hundreds of frames, the corresponding scattering amplitudes can be temporally averaged, resulting in a single, essentially noise-free scattering amplitude for each particle that is used for downstream analysis. In the following, we denote the measured scattering amplitude as |*A*(*q*)|, where *q* is the scattering vector defined as *q* = 2*k* sin(*θ/*2), with *θ* denoting the scattering angle.

**Figure 1:**
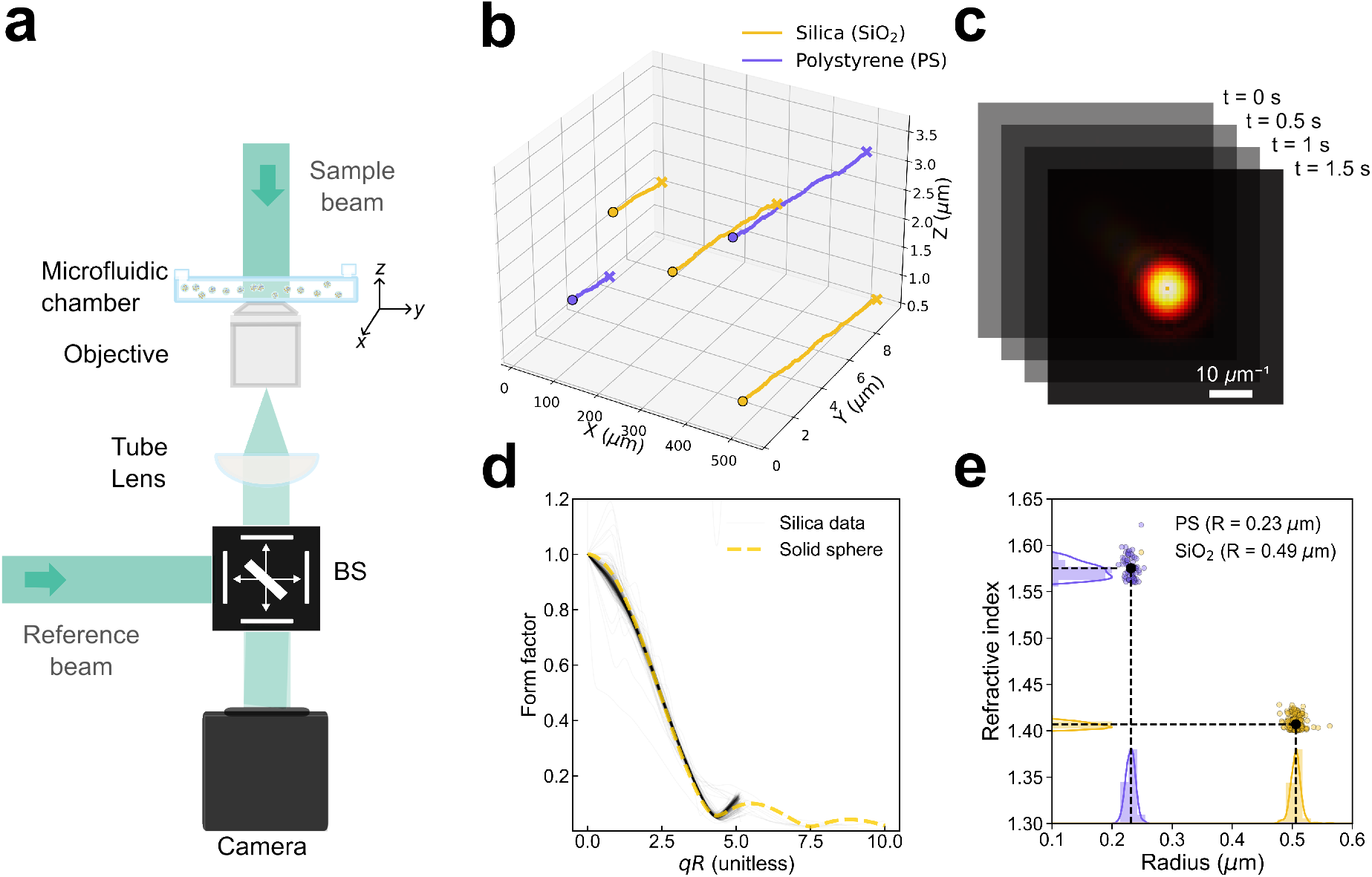
Quantitative analysis of holographic nanoparticle tracking data enables assessment of scattering form factor of individual submicrometer particles. (**a**) Schematic of the off-axis holographic microscopy setup using illumination at a wavelength of 532 nm. The illumination is split into a reference beam and a sample beam that passes through a microfluidic channel containing the sample. The scattered and transmitted light are collected by a 40× oil-immersion objective (NA 1.4), focused through a tube lens, and recombined with the reference beam at a beam splitter (BS) before reaching the camera. (**b**) Three-dimensional trajectories of individual nanoparticles obtained from holographic tracking within the microfluidic channel. Five representative traces are shown: two silica particles (SiO_2_, yellow) and three polystyrene particles (PS, purple). (**c**) Representative Fourier-transformed scattering patterns of individual tracked particles at selected time points (*t* = 0, 0.5, 1.0, and 1.5 s). Each panel shows the angular distribution of scattered light at the back focal plane. The scale bar corresponds to 10 µm^−1^ in spatial frequency. (**d**) Experimental scattering form factor as a function of *qR*, where *q* is the magnitude of the scattering wave vector and *R* is the particle radius. Solid black lines represent experimental data for silica particles, and the dashed yellow curve shows the theoretical Mie scattering prediction. The position of the experimental minimum matches the theoretical curve, confirming accurate size and refractive index retrieval. (**e**) Refractive index versus optical radius for the reference particles: polystyrene (purple, *R* = 0.23 µm, *n* = 1.58) and silica (yellow, *R* = 0.49 µm, *n* = 1.42). The distributions illustrate the measured variability within each population, with histograms displayed along the axes. This analysis pipeline enables simultaneous extraction of particle size and refractive index from single-particle holographic scattering measurements.

To gain intuition for how biophysical characteristics of submicrometer particles are encoded in these scattering amplitudes, we begin by considering the limit of small *q* (such that *qR* ≲ 3, where *R* is a characteristic particle size). In this regime, the scattering amplitude is approximated by an extended Guinier expression (see Supporting Information),

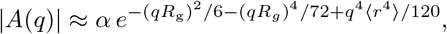

where *R*_g_ is the radius of gyration of the particle, and where we have defined

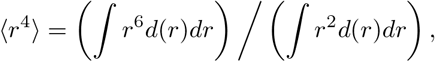

with *d*(*r*) denoting the local mass density of the particle at a radial distance *r* away from the particle center. For a solid sphere of radius *R, d*(*r*) is a constant within the radius and the above reduces to

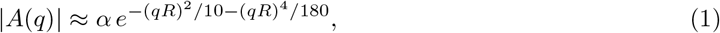

with the particle polarizability given by

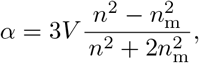

where *V* denotes the particle volume, and *n* and *n*_m_ are the refractive indices of the particle and the surrounding medium, respectively. This expression is an extension of the Guinier approximation commonly used in small-angle scattering [19] and highlights that the scattering amplitude encodes information about both particle composition, through the polarizability *α*, and particle size, through its dependence on *R*. Beyond the small-*qR* regime, the full scattering amplitude of a homogeneous sphere is described exactly by the Mie solution to Maxwell’s equations for electromagnetic radiation [20]. We define the optical radius *R*_opt_, hereafter *R*, as the particle radius which provides the best fit of Eq. (1) to the low *qR*-regime of the optical scattering pattern.

We validated the approach by fitting Eq. (1) to the individual scattering amplitudes of monodisperse silica spheres (nominal radius *R* = 0.49 µm provided by the supplier, *n* = 1.41) and polystyrene spheres (nominal radius *R* = 0.23 µm provided by the supplier, *n* = 1.58) measured in our system. While Eq. 1 captures only the small-angle (low-*q*) behavior, the full angular dependence of a homogeneous sphere is described by Mie theory. The measured scattering amplitudes closely follow the corresponding Mie solutions over the range of scattering angles accessible to the objective (Fig. 1**d** and SI Fig. 1), confirming that the reconstructed signal faithfully captures the angular distribution of the scattering amplitude for each particle. Moreover, the retrieved sizes and refractive indices for the individual particles in the monodisperse samples agree precisely with the nominal values (Fig. 1**e**).

### Measuring scattering patterns and protein concentration on a single condensate level

Having validated the approach on monodisperse solid particles, we next applied it to Ddx4N1, the N-terminal IDR of a DEAD-box helicase Ddx4, a widely used model system for biomolecular condensates [21–26]. We initially refer to the resulting Ddx4N1-rich objects as condensates; later structural analysis reveals that not all detected objects are consistent with compact phase-separated droplets.

A common feature of phase-separated systems is that the resulting distribution of condensate sizes is very broad [27–30]. Traditional characterization techniques either rely on ensemble measurements, such as bulk light scattering [17, 31, 32], or focus exclusively on large condensates that sediment to the bottom of the sample, thereby excluding the abundant population of microscopic condensates. Qualitative size distributions of microscopic condensates can be obtained by correlating fluorescence intensity to condensate size [25], but quantitative measurements are still lacking.

Using our approach, we track and analyze all condensates sufficiently large (*R* ≳ *λ/*2) to be accurately sized within the microfluidic device, while sufficiently small to avoid sedimentation (*R* ≲ 2 µm, enabling measurement of hundreds of individual condensates each minute. In contrast to the monodisperse reference particles, the scattering patterns of Ddx4N1 condensates formed at 10 *µM* protein, and 66 mM NaCl are highly heterogeneous (Fig. 2**a**, upper panel). An ensemble measurement would average over this variability and obscure the underlying polydispersity, as illustrated by the mean scattering amplitude shown as a dashed line in Fig. 2**a**. To estimate the sizes of the Ddx4N1 droplets, we fit the low-*q* region of each scattering amplitude using Eq. (1). The Guinier form provides a characteristic size *R* and the polarizability *α* = |*E*(0)|. After normalization by the estimated polarizability and rescaling by the fitted optical radius, the scattering amplitudes collapse in the low-*qR* regime onto the prediction for compact spheres (Fig. 2**a**, lower panel). This supports extraction of optical size and polarizability from the low-*q* signal. At larger scattering vectors, however, substantial particle-to-particle deviations become apparent, as will be examined in a later section.

**Figure 2:**
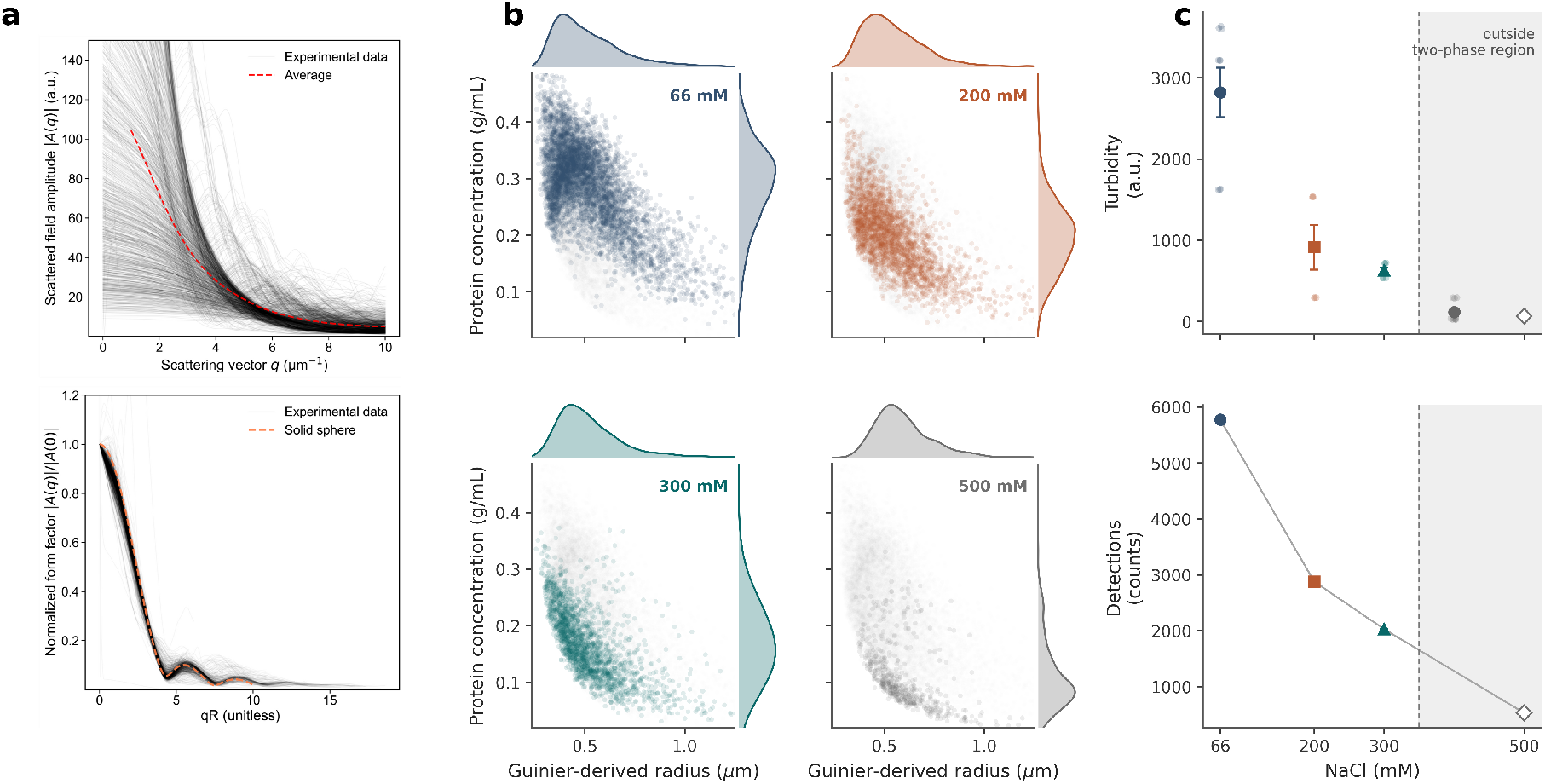
Holographic nanoparticle tracking enables high-throughput characterization of condensate physical properties. (**a**) Scattered-field amplitudes |*A*(*q*)| for all recorded condensates of Ddx4N1 (black curves, upper panel), illustrating the strong heterogeneity of the sample. The dashed red line shows the ensemble-averaged amplitude, corresponding to the type of signal that would be obtained in traditional ensemble small-angle scattering and highlighting how population-level averaging obscures single-condensate variability. The lower panel shows normalized scattering amplitudes |*A*(*q*)|*/*|*A*(0)| plotted as a function of *qR*, demonstrating that all single-particle curves collapse onto a common master curve for small values of *qR*. The experimental curves for Ddx4N1 condensates (black) are compared with the Mie-theory prediction for a homogeneous solid sphere (orange dashed line). (**b**) Optically inferred protein concentration versus optical radius for thousands of individual Ddx4N1 condensates, formed at 10µM protein concentration, measured at four NaCl concentrations (66 mM, 200 mM, 300 mM, 500 mM). Histograms indicate the distributions of sizes and mass concentrations under each condition. As salt increases, the refractive index systematically decreases, and the number of detectable condensates is reduced, consistent with suppressed phase separation at elevated ionic strength. (**c**) The turbidity of samples prepared at the same protein concentration as in (**b**) demonstrates that the three lowest salt concentrations lie inside the phase separating region, while the highest protein concentration lies outside the phase separating region. This is also reflected in a considerable reduction in detection count at the highest salt concentration.

For biomolecular solutions, the refractive index *n* reflects the internal mass concentration *c* through the refractive index increment (*dn/dc*) [33], as *c* = (*n* − *n*_med_)*/*(*dn/dc*), with *dn/dc* ≈ 0.19 mL*/*g being a typical value valid for most proteins [34]. This allows us to convert the measured condensate refractive index into an estimate of the mass concentration of protein inside the condensates. We find that while the estimated sizes of the condensates span a broad distribution, the optically inferred protein mass concentration (*c*_opt_) is centered around 0.31 g/mL (11 mM) within the condensates (Fig. 2**b**), consistent with previous findings [21,35], and suggesting a thousand-fold protein enrichment inside the condensate. The average *c*_opt_ decreases slightly with increasing condensate size. This effect is not seen in synthetic polymer systems, for which *c*_opt_ shows no dependence on size (SI Fig. 2).

The single-particle sensitivity of our approach enables quantitative analysis of how environmental parameters modulate condensate size and internal composition. Increasing salt weakens the electrostatic interactions that contribute to Ddx4N1 phase separation [21, 36, 37], which is expected to reduce condensate stability and decrease their internal mass concentration, and therefore their *c*_opt_. Consistent with this expectation, we observe a monotonic decrease in *c*_opt_ with increasing NaCl concentration (Fig. 2**b**). Coarse-grained molecular dynamics simulations of Ddx4N1 condensates recapitulate this trend, showing a decrease in dense-phase concentration with increasing ionic strength (SI Figs. 6–7). Turbidity measurements showed that the examined conditions span the turbidity-defined two-phase region, with the 500 mM NaCl condition lying outside the inferred phase boundary (Fig. 2**c**). Crossing the phase boundary was accompanied by a pronounced decrease in the detection count, although a residual population remained detectable at 500 mM NaCl (Fig. 2**c**).

We have validated the results presented in this section in a second, independently purified batch of Ddx4N1, as shown in SI Figs. 2, 4 and 5.

### Single-particle scattering reveals salt-dependent morphological heterogeneity

We next asked whether all detected Ddx4N1 assemblies are structurally consistent with compact condensates. To answer this question, we go beyond the Guinier approximation and quantify the scattering amplitude at *qR >* 1, which carries information about the assembly morphology. While the scattering amplitudes shown in Fig. 2**b** overlap closely in the Guinier regime (*qR <* 1), they begin to diverge near the first minimum of the scattering amplitude, which occurs around *qR* ≈ 4.1. This divergence reveals structural heterogeneity that would be obscured by ensemble averaging. Visual inspection of the scattering patterns measured by individual assemblies reveal two distinct morphologies (Fig. 3**a,b**). One population has a distinct, radially symmetric minimum and is therefore broadly consistent with compact spherical condensates. We refer to this population as sharp-interface condensates. The second population lack a well-defined minimum, despite still being approximately radially symmetric, and are referred to as the fuzzy-interface assemblies. Thus, assemblies with similar Guinier-derived sizes can nevertheless possess substantially different structures at larger scattering vectors.

**Figure 3:**
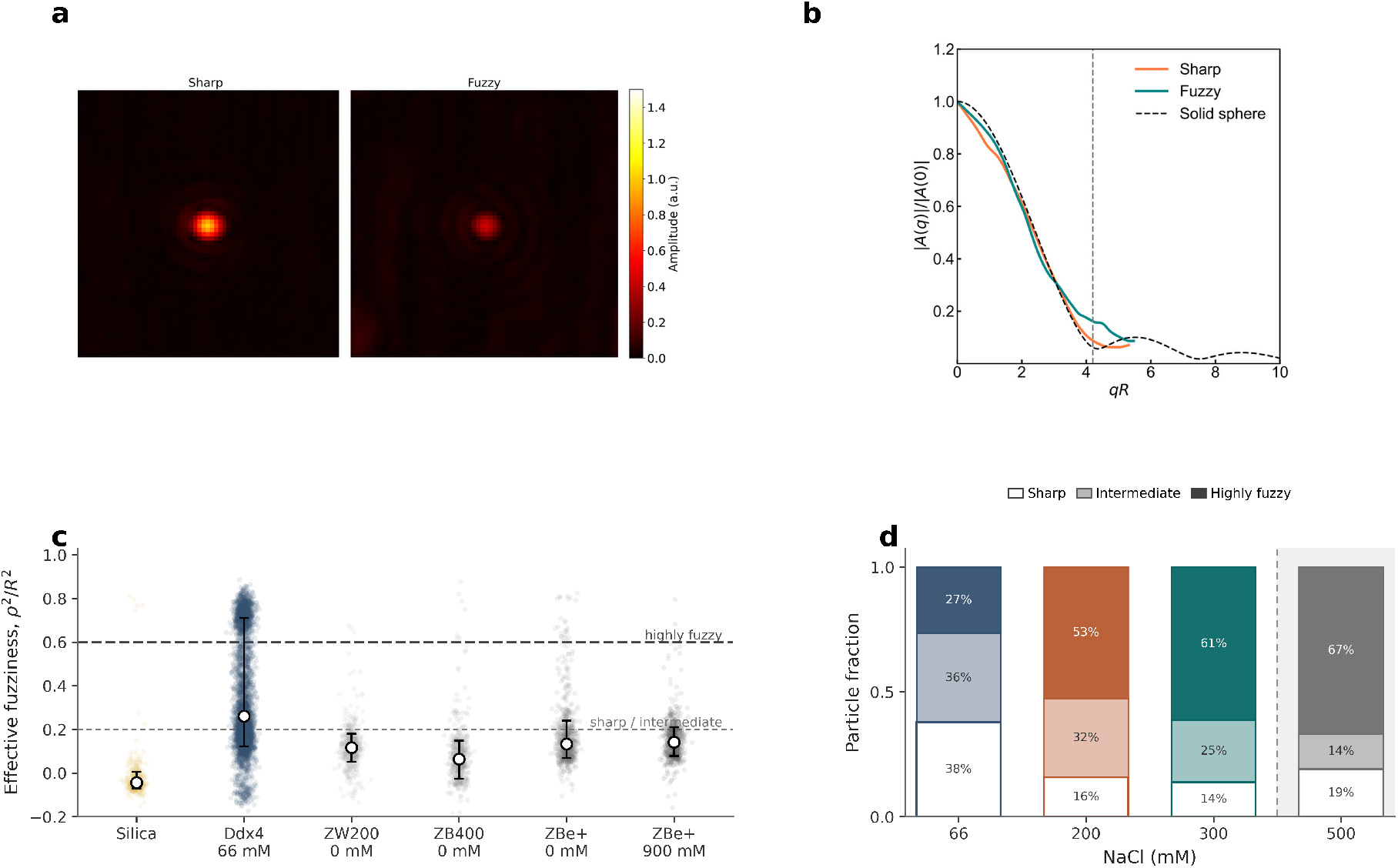
Single-particle holographic scattering reveals the emergence of a population of fuzzy assemblies in Ddx4N1 condensates. (**a**) Representative reconstructed scattering images of sharp and fuzzy Ddx4N1 assemblies of comparable size (*R* ~ 0.5 µm). (**b**) The corresponding normalized scattering amplitudes |*A*(*q*)|*/*|*A*(0)| as a function of the dimensionless scattering vector *qR*, compared with the theoretical scattering amplitude of a homogeneous solid sphere (black dashed line). The first minimum of the solid-sphere form factor (vertical dashed line) is suppressed for fuzzy assemblies, indicating the absence of a well-defined interface. (**c**) Distributions of the normalized interfacial-width parameter *ρ*^2^*/R*^2^ for Ddx4N1 condensates formed at 66 mM NaCl and for polymer condensates. The lower dashed line indicates the threshold used to classify assemblies as sharp, whereas the upper dashed line the threshold for classifying detections as highly fuzzy. Intermediate assemblies fall between those thresholds. (**d**) The fraction of assemblies with highly fuzzy interface increases while the fraction of assembly with sharp interfaces decreases with salt concentration.

To quantify this distinction, we fit the scattering amplitudes using a modified Mie scattering model that incorporates a fuzzy boundary through an effective interfacial-width parameter *ρ*^2^ (see Materials and Methods and Supplementary Information). We report the dimensionless quantity *ρ*^2^*/R*^2^, which normalizes this parameter by Guinier-derived assembly size. Small values correspond to scattering patterns with a pronounced minimum, corresponding to assemblies with a sharp and well-defined interface, while larger values describe progressive broadening or suppression of the minimum due to a more complex interfacial structure. Only assemblies with *R >* 0.4 µm were included in the detailed morphological analysis, as robust fitting of *ρ*^2^ requires scattering information beyond the low-*qR* Guinier regime.

To validate the extraction of this parameter, we first applied the model to homogeneous silica spheres with sharp, well-defined interfaces. As expected, their fitted values of *ρ*^2^*/R*^2^, were close to zero (Fig. 3**c**). By contrast, Ddx4N1 assemblies exhibited coexisting populations of sharp and fuzzy interfaces. To investigate whether this is a general feature of phase separating systems, we next examined phase-separated condensates formed by a series of controlled polymer systems containing electrostatic stickers (ZB400), electrostatic stickers with hydrophilic spacers (ZW200), or a combination of electrostatic and hydrophobic stickers with hydrophilic spacers (ZBe+). In these systems, fuzzy-interface assemblies were largely absent. ZBe+ exhibited a small fuzzy-assembly fraction at low salt concentration, but this population diminished when electrostatic interactions were screened by high salt (Fig. 3**c**). These controls show that a pronounced fuzzy population is not a generic consequence of liquid-liquid phase separation or electrostatic screening. Its greater prevalence in Ddx4N1 may therefore reflect the more complex, sequence-dependent interaction network of the protein.

At 66 mM NaCl, approximately 38% of the Ddx4N1 detections were classified as having a sharp interface, and 27% as having a highly fuzzy interface. The sharp interface fraction dropped to approximately 16% at higher salt, while the highly fuzzy fraction gradually increased in prominence, to 67% beyond the inferred phase boundary (Fig. 3**c,d**). The relative representation of sharp-interface condensates therefore decreased as the inferred phase boundary was approached, whereas a substantial population of fuzzy-interface assemblies remained among the sparse residual detections at 500 mM. Because the particles detected at 500 mM had low optical contrast and the total detection rate was strongly reduced (Fig. 2**c**), this fraction describes the composition of the detected and morphology-resolvable population rather than the absolute abundance of either morphology. Across the salt series, the relative representation of sharp-interface assemblies therefore decreased with increasing ionic strength, while fuzzy assemblies became progressively more prominent among the morphology-resolvable population. Coarse-grained molecular dynamics simulations of the condensed phase of Ddx4N1 showed that increasing salt altered the composition of the intermolecular contact network. Contacts between charged residues dominated at low salt, whereas aromatic contacts constituted a larger fraction of the remaining contacts at higher salt within the phase-separation regime (SI Fig. 3-5), providing a possible molecular correlate of the experimentally observed structural changes.

### Fuzzy assemblies are consistent with incomplete fusion

The fuzzy assembly profiles from Fig. 3**b** could arise either from spherical assemblies with diffuse boundaries, or from structurally heterogeneous assemblies. To distinguish these possibilities, we examined the scaling between mass and optical radius for the fuzzy assembly population. Compact assemblies of uniform density are expected to display *M* ∝ *R*^3^, whereas non-compact aggregates instead display 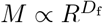, with the fractal dimension *D*_f_ *<* 3.

The fuzzy assemblies (*ρ*^2^*/R*^2^ *>* 0.2) consistently displayed a reduced scaling exponent (*D*_f_ ∼ 2.5), consistent with fractal organization (Fig. 4**a**), while sharp-interface assemblies displayed a scaling consistent with compact spheres (*D*_f_ ~ 3, SI Fig. 6). The fractal dimension was considerably reduced outside the two-phase region (*D*_f_ ~ 1.8 (Fig. 4**b**). The estimated fractal dimension outside the phase separating region is similar to theoretical estimates of the fractal dimensions of diffusion-limited cluster aggregation (*D*_f_ ~ ~ 1.6 − 1.8 [38, 39]), while the larger fractal dimensions observed inside the phase separating region suggests a more compact morphology, for instance due to partial fusion of the primary aggregation units [40].

**Figure 4:**
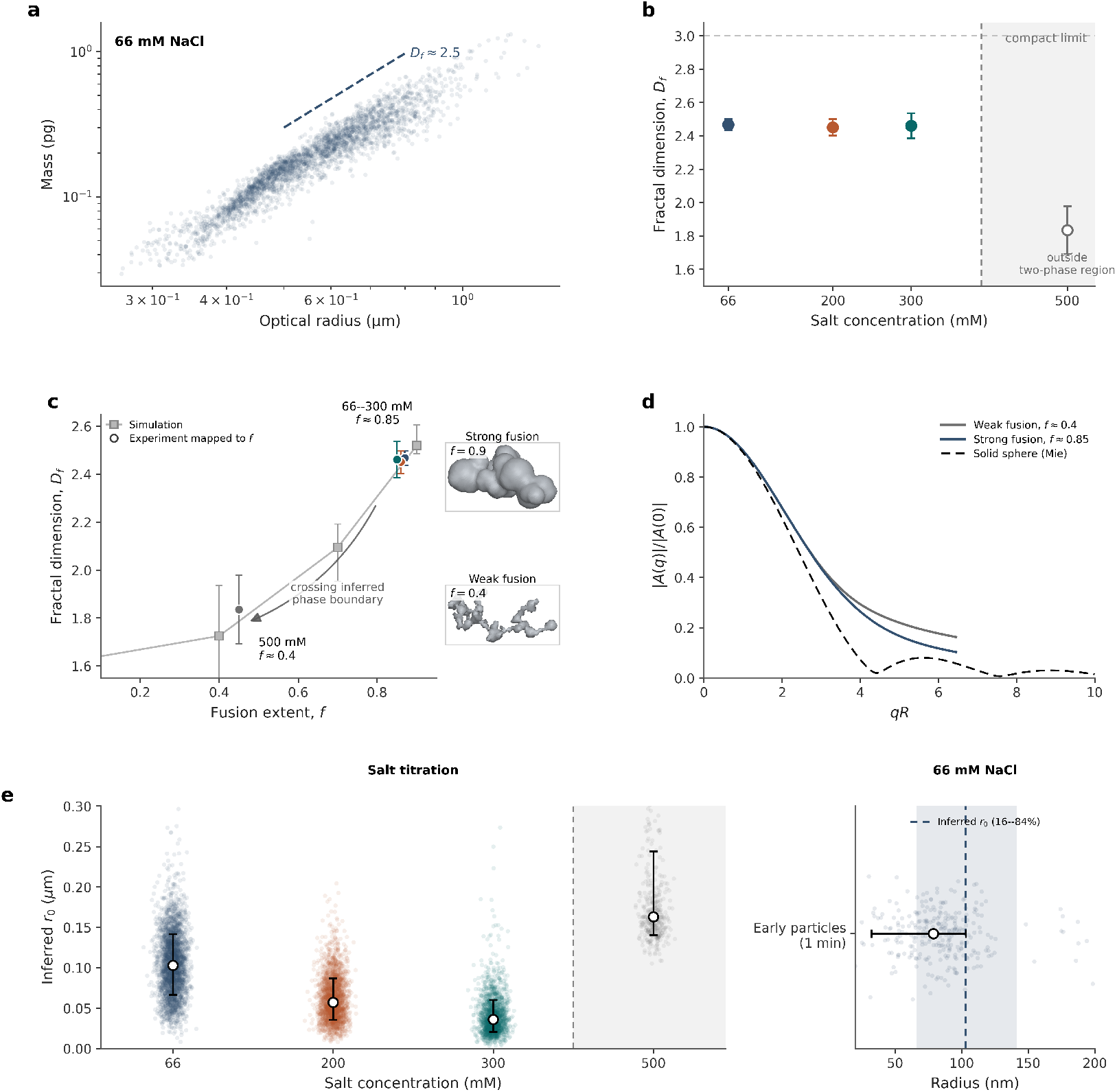
A partial-fusion model captures structural features of the fuzzy assemblies. (**a**) The population of Ddx4N1 fuzzy assemblies transition display a fractal scaling Mass ∝ *R*^2.5^. (**b**) The estimated fractal dimension for the fuzzy assembly population drops from *D*_f_ ~ 2.5 inside the phase separating region, to *D*_f_ ~ 1.8 outside the phase separating region (**c**) Experimental fractal dimensions map onto progressively less compact simulated aggregates with increasing salt. (**d**) Normalized scattering amplitudes for simulated partially fused aggregates lack a pronounced minimum and agree qualitatively with the experimental scattering amplitudes. (**e**) The inferred size of the primary units forming the aggregates decreases as the system approaches the inferred phase boundary. The inferred primary unit size are comparable in magnitude to a transient population of spherical particles observed using an orthogonal characterization approach.

To test this interpretation, we performed numerical simulations of diffusion limited cluster aggregation where the primary units were allowed to partially fuse upon contact. The degree of fusion was controlled by the fusion extent, *f*, of primary units, with *f* = 0 corresponding to rigid sticking and *f* = 1 to complete fusion. Based on these Brownian aggregation simulations, we found that the estimated fractal dimensions are consistent with aggregates formed with high degree of fusion inside the phase separating region (*f* ~ 0.85), and a low degree of fusion outside the phase separating region *f* ~ 0.4) (Fig. 4**c**). Moreover, the calculated scattering amplitudes of the simulated aggregates qualitatively reproduced the characteristic features observed experimentally (Fig. 3**d**), providing additional support for this interpretation.

We next used the aggregate scaling model to infer the characteristic radius *r*_0_ of the primary scattering units (Fig. 4**e**). At 66 mM NaCl, the inferred primary unit size was comparable to the radius of a transient population of small particles detected independently using DAISY-NTA [41] (Holtra AB), which can detect smaller/less dense particles, immediately after induction of phase separation. The abundance of these approximately 70-nm radius particles subsequently decreased. This agreement provides independent support for the inferred primary-unit scale and is consistent with, although it does not by itself prove, their participation in the formation of larger fractal assemblies.

### Optical and hydrodynamic radii reveal morphology-dependent hydrodynamics

We next investigated the hydrodynamic properties of the Ddx4N1 assemblies. Individual assemblies were tracked over several hundred frames as they moved through the microfluidic channel, providing access to both their scattering amplitudes and their Brownian motion. By combining trajectories with sizes extracted from the corresponding scattering patterns, we directly compared the optical radius, *R*_opt_, with the hydrodynamic radius, *R*_h_, inferred from diffusivity.

The hydrodynamic radius is defined through the Stokes relation,

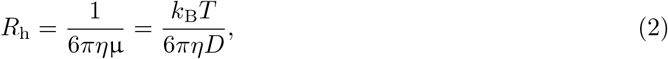

where *η* is the solvent viscosity, µ is the hydrodynamic mobility, and *D* is the translational diffusion coefficient. The diffusion coefficient was extracted from the statistics of individual trajectories (Materials and Methods). To obtain quantitative estimates of *R*_h_, the viscosities of all buffer conditions were measured independently (Materials and Methods).

Whereas the Guinier-derived optical radius characterizes the spatial extent of the scattering mass, the hydrodynamic radius measures the drag exerted by the particle on the surrounding fluid. More precisely, *R*_h_ is the radius of an equivalent no-slip sphere with the same translational drag. For a rigid spherical particle with a no-slip boundary, the hydrodynamic and physical radii coincide. For soft particles such as condensates, internal circulation, interfacial mobility, surface fluctuations, and solvent penetration can reduce the drag, such that the effective hydrodynamic boundary does not coincide with the optically defined boundary (Fig. 5a).

**Figure 5:**
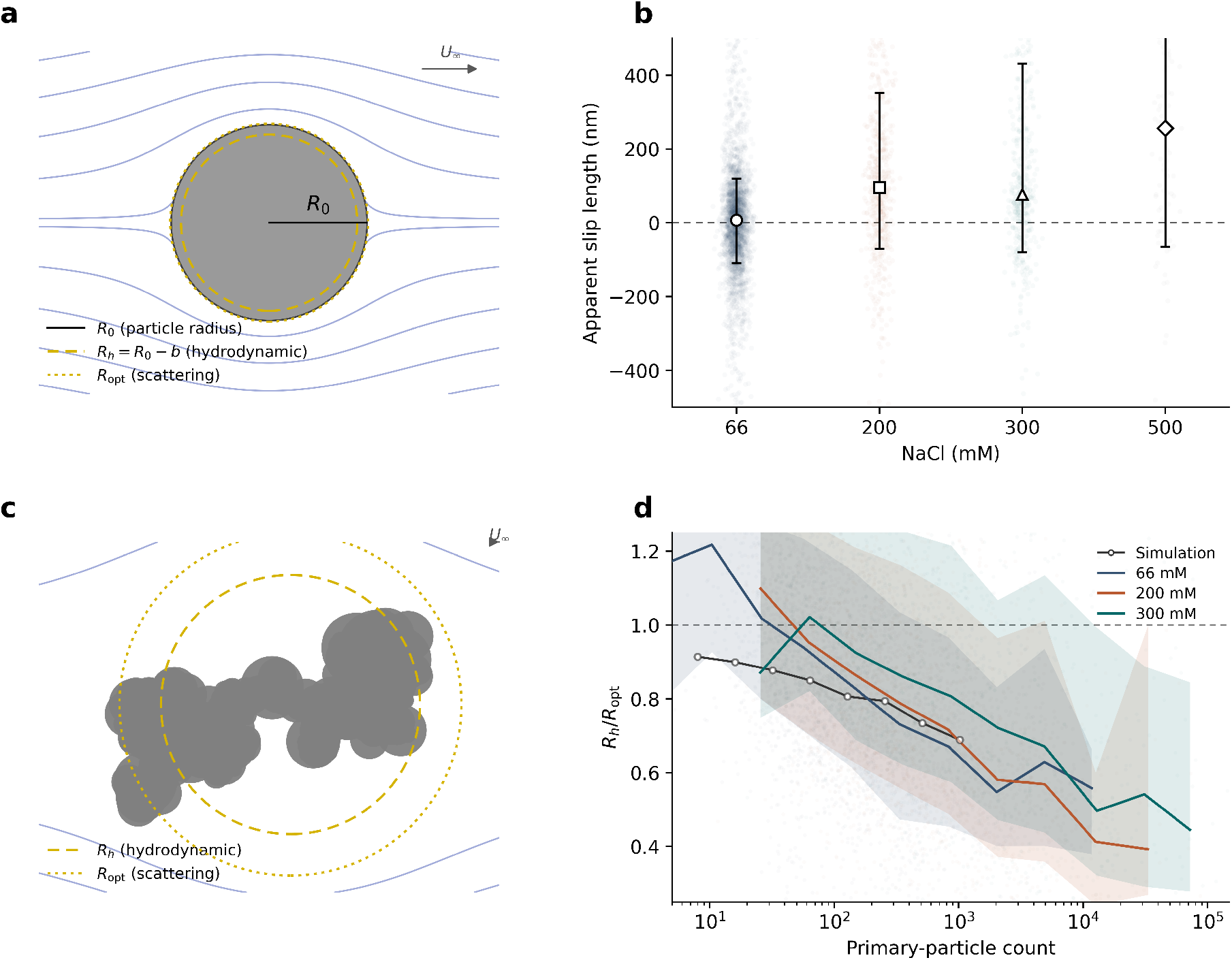
Hydrodynamic radius acts as a probe for interfacial physics. **(a)** The hydrodynamic radius essentially encodes the position at which the motion of the external fluid vanishes under Stokes flow. For a sharp interface, slip at the interface can cause the effective hydrodynamic boundary to be shifted toward the particle center. Meanwhile, the optical radius measures the distribution of mass within the particle, and is physically distinct from the hydrodynamic radius. **(b)** Condensates with a sharp boundary display a lower hydrodynamic radius than the optical radius, suggesting interfacial slip. **(c)** For an aggregate, the hydrodynamic radius is typically smaller than the optical radius due to aggregate permeability. **(d)** The ratio between hydrodynamic radius and optical radius decreases as a function of the number of monomers in the measured aggregates, in agreement with Brownian aggregation simulation results.

We therefore define an apparent slip length,

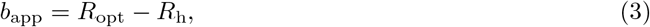

which quantifies the displacement between the optically defined boundary and the radius of the equivalent no-slip sphere.

Within the sharp-interface population, *R*_h_ was generally smaller than *R*_opt_, corresponding to a positive *b*_app_. At fixed protein concentration, the apparent slip length increased with salt concentration as the system approached the inferred phase boundary (Fig. 5b). The salt dependence of *b*_app_ therefore indicates condition-dependent coupling between the optical structure of the particles and their hydrodynamic boundary, potentially reflecting changes in interfacial mobility, internal circulation, or solvent penetration. A positive apparent slip length was also observed in the polymer control systems, whereas monodisperse polystyrene particles yielded *b*_app_ ≈ 0, indicating that the discrepancy between optical and hydrodynamic radius is characteristic of the soft condensate systems examined here rather than a systematic measurement offset (SI Fig. 7).

For the fuzzy population, the hydrodynamic radius is additionally sensitive to the open and potentially solvent-permeable structure of the assembly(Fig. 5c). The ratio *R*_h_*/R*_opt_ decreased with increasing number of primary units in the aggregate, and the same trend was reproduced by the Brownian aggregation simulation (Fig. 5d). This behavior is consistent with increasingly open or hydrodynamically permeable structures at larger aggregate sizes, for which the increase in hydrodynamic drag is slower than the increase in optical size.

## Discussion

We have introduced interferometric imaging for multiparametric characterization of submicrometer biomolecular condensates with high throughput, analyzing hundreds of condensates per minute. This approach combines off-axis holographic microscopy with quantitative scattering analysis, enabling simultaneous determination of condensate size, refractive index, internal mass concentration, interfacial structure, and hydrodynamic mobility at the single-particle level. This technique fills an important gap in the current arsenal of condensate characterization techniques, as it allows to resolve structural and compositional heterogeneity across condensates in the size range 200 nm *< R <* 1500 nm. Based on the analysis of more than 15 000 individual Ddx4N1 assemblies, this combination of observables reveals that a chemically homogeneous condensate-forming system can populate distinct mesoscale architectures with different interfacial and hydrodynamic properties. Crucially, these populations are not distinguished simply by size and would therefore be obscured by ensemble-averaged measurements or methods that report only a single particle parameter.

Central to our approach is characterization through analysis of single particle scattering profiles. In the case of Ddx4N1, these profiles revealed two coexisting subpopulations that would be compressed into a single broad distribution by monoparametric techniques. Assemblies displaying a sharp minimum in the scattering profile are consistent with spherical condensates with a well-defined interface. The second population displayed a less sharp minimum, or lacked a minimum altogether. This is broadly consistent with a more extended, fuzzy interface. The relative prevalence of these structural phenotypes changes systematically across the salt series.

The simultaneous measurement of size and mass of individual assemblies offered by our approach further distinguishes spherical particles from non-compact assemblies. The mass–size scaling establishes that the larger fuzzy assemblies are non-compact, while Brownian aggregation simulations show that partial fusion of smaller protein-rich units provides a minimal physical model capable of reproducing both the observed scaling and the suppression of the scattering minimum. The present measurements do not, however, directly resolve the formation pathway. We therefore cannot exclude alternative mechanisms in which structural heterogeneity develops within initially compact condensates. Direct time-resolved observation of individual assembly and coalescence events will be required to uniquely identify the pathway.

The controlled polymer systems further illustrate the capacity of the method to connect molecular interaction design with mesoscale assembly structure. Two morphological populations were observed only when multiple sticker types were present, hinting that interaction heterogeneity, rather than multivalency alone, promotes competing assembly outcomes.

Finally, tracking the same assembly over hundreds of frames adds an independent hydrodynamic dimension to the characterization. We find that condensates display a condition-dependent reduction in hydrodynamic drag, indicating that the interfacial structure and coupling to the surrounding fluid changes across the salt series.

It is interesting to note, that the structural heterogeneity observed in the Ddx4 system is not captured by mean-field descriptions of liquid-liquid phase separation, in which the dense phase concentration and interfacial structure is determined completely by the coarse-grained interaction parameter governing the free energy functional. Instead, our results indicate that mesoscale structural heterogeneity can arise from molecularly identical systems. Amino-acid sequence, conformational fluctuations, and the spatial arrangement of interaction motifs may together establish a hierarchy of interactions that is apparent at submicrometer length scales but becomes obscured upon macroscopic or ensemble averaging. This highlights the need for high-throughput, multiparametric single condensate characterization to study such dynamic heterogeneity in condensate systems. In particular, we anticipate that a mechanistic understanding of how protein conformations and sequences influence such structural heterogeneity may help address several open challenges in the field, including understanding the pathways involved in condensate ageing [42], gelation [5], and condensate-mediated aggregation and amyloid formation [43], as well as the role of posttranslational modifications in dynamically regulating condensate assembly and disassembly [4]. More broadly, it provides a route to connect molecular interaction design to the emerging physical state of individual biomolecular assemblies.

## Materials and Methods

### Protein expression and purification

The N-terminal domain of Human DEAD-box helicase-4 (Ddx4; 1–236) with an N-terminal 6xHis tag was expressed in E. Coli BL21 Gold (DE3) cells. Briefly, the cells were transformed via heat shock at 42°C for 30 s and grown overnight at 37°C in Luria Broth agar plates supplemented with 100 µg ampicillin ml^−^1. Cultures were scaled up in rich media, and protein expression was induced by adding 0.5 mM IPTG, followed by incubation overnight at 20°C. Cells were harvested by centrifugation, resuspended in lysis buffer (1 M NaCl, 50 mM Tris at pH 8.5, 10 mM imidazole, 2 mM *β*-mercapthoethanol), and stored at –20°C.

The protein was first purified by affinity chromatography using a Ni^2+^-charged Fast Flow Chelating Sepharose matrix (Cytiva Sweden AB, Uppsala, Sweden) and by size exclusion with a HiLoad 16/600 Superdex 75 pg column (Cytiva Sweden AB) equilibrated in a buffer containing 50 mM TRIS pH 8.5, 1 M NaCl, and 10% glycerol. Selected fractions from size-exclusion chromatography were concentrated to ~150 µM using Amicon 10,000 MWCO spin filters, flash-frozen in liquid nitrogen in 10 µL aliquots, and stored at −80°C.

### Polymer synthesis

All polymers were synthesized via RAFT polymerization using 4,4-azobis(4-cyanovaleric acid) (ACVA) as an initiator and 4-cyano-4-(phenylcarbonothioylthio)pentanoic acid (CPA) as a RAFT agent according to Capasso Palmiero et al. [44]. The monomer concentration was set to around 10 wt %, and the initiator to CPA molar ratio was set to 1/3. ZW200, where 200 refers to the degree of polymerization contained 60 sulfabetaine methacrylate (ZB) monomers acting as stickers and 140 sulfobetaine methacrylate (SB) monomers acting as spacers. ZB400 contained ZB stickers only with a degree of polymerization of 400. ZBe+ contained 50 ZB stickers, 50 benzyl groups as stickers and 100 SB groups as spacers, and the degree of polymerization was 200.

### Sample preparation

Polystyrene (PS) particles with a nominal diameter of 456 nm (Thermo Fisher Scientific) were used as reference particles for point-spread-function (PSF) calibration and refractive-index validation. Silica particles with a nominal diameter of 985 nm (Thermo Fisher Scientific) were also measured as an additional solid-sphere reference.

In the case of Ddx4, phase separation was triggered by diluting the protein stock solutions in experimental buffer containing 20 mM TRIS at pH 7.5, containing the desired amount of NaCl, taking into account the salt concentration from the storage buffer. In the case of polymers, phase separation was triggered by diluting the stock solutions in MQ water containing the desired amount of NaCl.

### Use of AI tools

AI tools (ChatGPT, Claude) have been used to suggest improvements in visual and textual clarity, and for code generation/debugging. All suggestions were carefully reviewed and validated by the authors, and the authors take full responsibility for the content of the manuscript.

All mixtures were gently mixed by pipetting up and down after addition of NaCl and protein to ensure thorough mixing.

The polymer stock solution was prepared at 40 g L^−1^ in 1.2 M NaCl. For each measurement, a total volume of 500 µL was prepared using Milli-Q water and 3 M MgCl_2_ stock to set the desired salt concentration: All solutions were gently mixed by pipetting up and down after addition of MgCl_2_ and polymer to ensure thorough mixing. The samples were immediately introduced into the microfluidic channel for holographic imaging.

All measurements were performed at room temperature (25 °C).

### Brownian aggregation simulation

Formation of aggregates from partially fusing monomers was simulated using a Monte Carlo simulation of diffusion limited cluster-cluster aggregation using a Smoulochowski pair-selection kernel. The simulation starts by defining a set of monomers with a configurable polydispersity. Two existing clusters are chosen according to the pair selection kernel, and these are subsequently merged by performing a Brownian dynamics simulation of the motion of the clusters until they come into contact. Upon contact, the clusters partially fuse according to a configurable fusion extent parameter, and the two clusters merge into a single, larger cluster. This process is repeated until only a single cluster remains.

The hydrodynamic radius is calculated by first constructing the Rotne-Prager-Yamakawa mobility tensor, imposing a common translational velocity on all beads, and subsequently rotationally average the hydrodynamic mobility.

### Molecular dynamics simulations

Simulations were performed using the CALVADOS 2 model [45, 46] implemented in OpenMM (v8.2) [47, 48]. 200 chains were simulated in a box of dimensions [20, 20, 300] nm at 298.15 K using a Langevin integrator with a time step of 10 fs and a friction coefficient of 0.01 ps^−1^. The charge of the His residues was set to 0.03 *e* based on the Henderson-Hasselbalch equation (pH=7.5, p*K*_*a*_ = 6). Simulations were carried out for 11 µs, and the first 200 ns were discarded for equilibration. The effect of salt concentration on charge-charge interactions was modeled through the Debye-Hückel potential with dielectric constant *ϵ*_*r*_ = 78.4 and ionic strength 66, 216, and 366 mM. All the simulated systems formed a protein-rich slab coexisting with a dilute protein solution. To estimate the thickness of the interface, *d*, we fitted the two halves of the concentration profiles to the following function

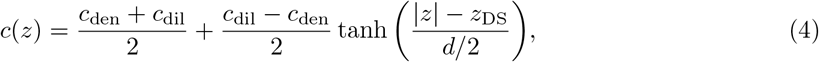

where *c*_den_ and *c*_dil_ are the average concentrations of the dense and dilute phases, and *z*_DS_ is the position of the dividing surface [47]. Intermolecular contacts between a chain in the middle of the slab and the surrounding chains were calculated as

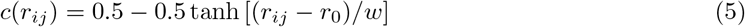

where *r*_*ij*_ is the residue-residue distance, *r*_0_ = 1 nm, and *w* = 0.3 nm [47].

### DAISY characterization

Data was acquired and analyzed using DAISY-NTA (Holtra AB, Gothenburg, Sweden). The instrument performs Nanoparticle Tracking Analysis (NTA) using Dual Angle Interferometric Scattering microscopY (DAISY) and holographic imaging. The instrument detects both backscattered and forward-scattered light, enabling size and refractive index determination over a size range of about 50-1000nm diameter (silica particles).

Similar microfluidic chips as descibed below for the main method were used, pre-incubated with BSA. A series of 5 s recordings, covering a field of view of 45×45 mm, were made of the sample under flow, for both imaging modalities at 800fps. From mixing of buffer and protein sample, the delay time to reach field of view and starting recording was about 60-90s. Three such measurements were made.

### Microfluidic channel preparation

Measurements were performed in a ChipShop^®^ microfluidic channel fabricated from Topas with a depth of 20 µm, a width of 800 µm, and a total channel length of 58.5 mm. The sample solution was loaded into the channel inlet, and measurements were carried out under continuous flow conditions.

Before sample loading, a single channel of the microfluidic chip was pretreated with bovine serum albumin (BSA, 5%) to prevent adhesion of condensates to the channel walls. The channel was incubated with the BSA solution for 3 minutes and subsequently rinsed several times with Milli-Q (MQ) water. This surface treatment produced a uniform, hydrophilic coating that minimized nonspecific binding of condensates and particles to the channel surface.

In microfluidic flow, the relevant regime is laminar flow, where streamlines are parallel and mixing occurs primarily through diffusion.^1^

### Imaging and data acquisition

Measurements were performed using an off-axis holographic microscope equipped with a 532.141 nm continuous-wave laser (coherence length *>* 1 m; model MGL-III-532 nm-150 mW-EL2163) and a 40× oil-immersion objective (NA 1.4). The sample and reference beams were recombined at a beam splitter and imaged onto a high-speed CMOS camera (Basler acA1300-200um) operating at 270 frames s^−1^. Each measurement comprised three independent videos; each recording lasted 40 s, corresponding to 10,800 frames at 270 frames s^−1^ (40 s × 270 s^−1^). The typical field of view contained ~30 particles. Across replicates, hundreds of single-particle trajectories were analyzed per condition.

### Data processing and analysis

Single-particle tracking was performed in MATLAB, using analysis routines previously applied in related work [12, 41].

The system point-spread function (PSF) was obtained from reference polystyrene particles by averaging the reconstructed complex field over time and taking the absolute value of the mean amplitude, yielding a real, positive scattering envelope. This PSF was used to normalize experimental scattering patterns and correct for the instrument response before quantitative analysis.

For each particle trajectory, the diffusion constant was estimated using a covariance-based estimator [49]. The hydrodynamic radius *R*_*h*_ was then calculated using the Stokes–Einstein relation,

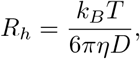

where *k*_*B*_ is the Boltzmann constant, *T* = 300 K, and *η* is the buffer viscosity. Viscosities were measured independently using a viscometer for protein- and polymer-free buffer solutions: 1.044, 1.054, and 1.208 mPa·s for Tris buffer at 200, 300, and 500 mM NaCl, respectively, and 1.079, 1.122, and 1.208 mPa s for water with MgCl_2_ at the same concentrations. The appropriate viscosity was used for each Stokes–Einstein calculation.

For optical characterization, the Fourier transform of each particle’s reconstructed field yielded its angular scattering amplitude *A*(*q*). At low scattering vectors, fitting was performed using the Guinier approximation (Eq. 1), where |*A*(*q*)| encodes both the particle polarizability and its radius of gyration. At higher *qR*, normalized scattering amplitudes were compared with full Mie theory for homogeneous spheres to extract the optical radius (*R*_opt_), refractive index (*n*_*p*_), and interfacial-fuzziness parameter (*ρ*^2^). Reported uncertainties correspond to the standard deviation across all analyzed particles within each condition. All subsequent data fitting and statistical analyses were performed in Python using custom scripts.

## Code and data availability

Code needed to reproduce the molecular dynamics simulation results is available at https://github.com/gitesei/_2026_Ddx4N_simulations. All data is available from the authors upon request.

## Acknowledgements

The authors thank Roberto Frigerio, Suiying Ye and Niccolo Zanieri for providing polymer samples. D.M. acknowledges funding from Olle Engkvist foundation and the Swedish Research Council (grant number 2019-05071). P.A and K.M. acknowledges the European Research Council through the Horizon 2020 research and innovation programme (grant agreement No. 101002094, P.A.) for financial support. G.T. acknowledges a donation to Water Science Lab from Sandberg Development. Molecular dynamics simulations were enabled by resources provided by LUNARC, The Centre for Scientific and Technical Computing at Lund University.

## Supplementary information

This section provides detailed information about the materials and experimental procedures employed in this study.

## Footnotes

1 In laminar flow, no turbulent mixing occurs; this is characteristic of the low Reynolds numbers typically found in microfluidic environments.

