## Supporting information for "Single-particle light scattering reveals structural heterogeneity of biomolecular condensates"

Berenice García Rodríguez[1], Katarzyna Makasewicz[2], Giulio Tesei[4,5], Fredrik Eklund[6], Edvin Johansson[6], Paolo Arosio[2], Giovanni Volpe[1,3], and Daniel Sundås Midtvedt[1]

[1]Department of Physics, University of Gothenburg, Gothenburg, Sweden
[2]Department of Chemistry and Applied Biosciences, ETH Zurich, Zürich, Switzerland
[3]Science for Life Laboratory, Department of Physics, University of Gothenburg, Gothenburg, Sweden
[4]Department of Biomedical Science, Malmö University, Malmö, Sweden
[5]Biofilms Research Centre for Biointerfaces, Malmö University, Malmö, Sweden
[6]HOLTRA AB, Sahlgrenska Science Park, Gothenburg, Sweden

September 10, 2026

### 1 Description of the analysis of individual scattering patterns

#### 1.1 Preprocessing of the data

For each detected particle, a small region of interest (ROI) centered on the particle was extracted from the numerically refocused hologram. The complex optical field within this ROI was obtained from the off-axis holographic reconstruction and background-subtracted to remove static interference contributions. Since each particle was detected multiple times, each particle is represented by a stack of numerically refocused and centered ROIs. Prior to further analysis, this stack was temporally averaged to reduce noise. The angular scattering amplitude was computed by first applying a two-dimensional Fourier transform to the complex field in the ROI. In Fourier space, the radial coordinate is related to the angle $\theta'$ at which the scattered light enters the objective as [1]

$$k_{\parallel} = \frac{2\pi}{\lambda} \sin(\theta'),$$

where $\lambda$ is the illumination wavelength. Since the immersion medium ($n'_m = 1.515$) differs from the condensate medium ($n_m \approx 1.33$), the angle $\theta'$ is related to the scattering angle $\theta$ by Snell's law,

$$n'_m \sin(\theta') = n_m \sin(\theta).$$

The scattering wave vector $q$ is in turn related to the scattering angle $\theta$ by

$$q = \frac{4\pi}{\lambda} n_m \sin(\theta/2),$$

demonstrating how the amplitude how the Fourier-transformed field $|E(k_{\parallel})|$ is related to the scattering amplitude $|E(q)|$ through a simple coordinate transformation.

#### 1.2 Correction for motion blur

Since the particles are monitored under flow, the particles will move slightly during the exposure time of the camera. This will blur out the scattering pattern slightly in the direction of flow. Assuming linear motion in the x-direction, the measured scattering pattering will be related to the true scattering pattern by

$$E_{\text{meas}}(\vec{k}_{\parallel}) = \int_{-\Delta x/2}^{\Delta x/2} E(\vec{k}_{\parallel}) e^{ik_x \cdot x} dx = E(\vec{k}_{\parallel}) \frac{\sin(k_x \Delta x/2)}{k_x \Delta x/2},$$

where $\Delta x = v_{\text{flow}}\tau$, and where $v_{\text{flow}}$ is the flow speed and $\tau$ is the exposure time. We found that an exposure time of $\tau = 2$ ms seemed to correct for the resulting motion blur better than the experimentally set exposure time of 3.8 ms (making the patterns radially symmetric), so in the analysis an exposure time of 2 ms was used for correcting the motion blur.

##### 1.3 Quantification of the point spread function

In addition to motion blur, the scattering patterns will also be affected by the point spread function of the microscope. Specifically, in the back focal plane, the point spread function (PSF) will act as a multiplicative mask, so that

$$E_{\text{meas}}(\vec{q}) = \text{PSF}(\vec{q})E(\vec{q}).$$

To quantify the point spread function, a set of monodisperse polystyrene particles (nominal diameter 450 nm and refractive index 1.58, from Microparticles GmBH) was measured. The measured scattering patterns were compared to the numerically predicted scattering pattern of such microparticles using Mie theory calculations (ref to miepython), and the point spread function was estimated as

$$\text{PSF}(\vec{q}) = E_{\text{meas}}(\vec{q})/E(\vec{q})$$

.

##### 1.4 Derivation of the extended Guinier approximation

At small scattering angles such that $qR < 1$, with $q = (4\pi/\lambda)\, n_{\text{m}} \sin(\theta/2)$, and assuming $\Delta n \equiv n_{\text{p}} - n_{\text{m}} \ll n_{\text{m}}$, the scattering is determined by interference effects between the unperturbed external field and the scattered field. In this regime, the scattering amplitude is conveniently expressed as a Fourier transform of the refractive index distribution of the particle [1],

$$E(\vec{q}) = k\alpha \frac{\int d\vec{x}\Delta n(\vec{x})e^{i\vec{q}\cdot\vec{x}}}{\int d\vec{x}\Delta n(\vec{x})}$$

, in which $\alpha = 3V\left(n_{\text{p}}^2 - n_{\text{m}}^2\right)/\left(n_{\text{p}}^2 + 2n_{\text{m}}^2\right)$ is the particle polarizability. First of all, since the refractive index contrast is proportional to the mass concentration within the particle [2], the integral is essentially a Fourier transform of the mass distribution. Further, since we are working with small-angle scattering, it is safe to Taylor expand the exponential. Noting that for a spherically symmetric particle, odd terms vanish by symmetry, and keeping terms up to fourth order, the scattering amplitude can be written as

$$E(q) = E(0)\left(1 - q^2\langle r^2\rangle/6 - q^4\langle r^4\rangle/120\right), \tag{1}$$

where the brackets denote

$$\langle y\rangle = \frac{\int y\Delta n(\vec{x})}{\int \Delta n(\vec{x})}.$$

In this form, $\langle r^2\rangle \equiv R_{\text{g}}^2$ by definition, with $R_{\text{g}}$ being the radius of gyration, and $\langle r^4\rangle = (3/7)R_{\text{g}}^4$ for a sphere. Taking the logarithm of Eq. (1) and Taylor expanding the result yields an exponential approximation for the scattering amplitude,

$$|E(q)| \approx |E(0)| \exp\left[-\frac{(qR_{\text{g}})^2}{6} - \frac{(qR_{\text{g}})^4}{72} + \frac{q^4\langle r^4\rangle}{120}\right].$$

This is an extension of the regular Guinier approximation, keeping terms up to $(qR_{\text{g}})^4$. For a sphere, the extended Guinier approximation can be explicitly calculated in terms of the sphere radius $R$ as

$$|E_{\text{sphere}}(q;R)| \approx |E(0)| \exp\left[-\frac{(qR)^2}{10} - \frac{(qR)^4}{500}\right]. \tag{2}$$

By numerical simulations, we validated that this approximation remains valid for scattering vectors satisfying $qR < 3$. To model the scattering from a fuzzy sphere, we write the refractive index distribution as a convolution of that of a perfect sphere and a "fuzziness kernel" $\kappa$ as $\Delta n(\vec{x}) = \delta n(\Theta(\vec{x};R) * \kappa(\vec{x}))$, where $\delta n$ is the refractive index contrast in the homogeneous center of the fuzzy sphere, and $\Theta(\vec{x};R) = 1$

for $|\vec{x}| < R$ and 0 otherwise. Since the scattering amplitude is given by the Fourier transform of the refractive index distribution, one obtains

$$|E_{\text{fuzzy}}(q)| \approx |E(0)||E_{\text{sphere}}(q;R)\hat{\kappa}(q)|,$$

where $\hat{\kappa}(q)$ is the Fourier transform of the fuzziness kernel $\kappa(\vec{x})$. Taking, for simplicity, a Gaussian fuzziness kernel, one obtains

$$|E_{\text{fuzzy}}(q)| \approx |E(0)||E_{\text{sphere}}(q;R)| \exp\left[-\frac{q^2\rho^2}{2}\right]$$

where $\rho$ is the width of the Gaussian kernel. This expression shows that the effect of the fuzziness is to alter the behavior of the scattering amplitude at small scattering angles by a renormalization of the radius of gyration as $R_{\text{g}}^2 = R_{\text{g, sphere}}^2 + 3\rho^2$, while higher order terms in the scattering amplitude are determined completely by the radius of the homogeneous core, $R$. Thus, an even better approximation for the scattering from a fuzzy sphere can be obtained by replacing the extended Guinier approximation for the scattering from the homogeneous core by the exact scattering from a homogeneous sphere,

$$|E_{\text{fuzzy}}(q)| \approx |E(0)||E_{\text{Mie}}(q;R)| \exp\left[-\frac{q^2\rho^2}{2}\right], \tag{3}$$

where $E_{\text{Mie}}(q;R)$ is the Mie solution of the scattering from a homogeneous sphere. Within this approximation, the curvature of the scattering at low values of $q$ defines the radius of gyration of the particle, while the position of the first minimum in the scattering amplitude defines the radius of the homogeneous core.

#### 1.5 Estimation of size, refractive index, and fuzziness using the extended Guinier approximation

The estimation of size and refractive index of individual particles was performed by a multi-stage fitting procedure. First, the scattering amplitude $|E(q)|$ measured at all accessible scattering vectors $q$ was used to produce an initial estimate of $|E(0)|$ and $R$ using Eq. (2). Next, this initial size estimate was used to determine an approximate range of validity of the extended Guinier approximation. Specifically, scattering vectors satisfying $q > 0.5\mu\text{m}^{-1}$ and $qR < 3$ were used to perform a refined fit only in the extended Guinier region to refine the estimates of $|E(0)|$ and size $R$. Next, $|E(0)|$ was converted to a refractive index by comparing it to numerical simulations of the scattering amplitude at $q = 0$ using the exact Mie solution for a particle of size $R$, evaluated using the package MiePython [3]. Finally, to estimate the interface width, we fitted Eq. (3) to the scattering amplitude using values of the scattering vector satisfying $qR > 2$ and $qR < 6$ to focus on the behavior around the first minimum, and fixing the radius of gyration to that obtained in the first round of fitting.

#### 1.6 Mesoscopic simulations of aggregation with incomplete fusion

To investigate whether aggregation followed by incomplete fusion can account for the structural properties of the fuzzy Ddx4N1 assemblies, we implemented an off-lattice three-dimensional cluster–cluster aggregation model, following the general framework cluster-cluster aggregation models [4]. The model is intended as a mesoscopic description of aggregate geometry and does not explicitly simulate molecular interactions, fluid flow, or interfacial dynamics. Primary protein-rich units are represented as spherical mass elements of uniform density, such that the mass of element $i$ is proportional to

$$m_i \propto a_i^3, \tag{4}$$

where $a_i$ denotes its radius.

##### 1.6.1 Primary-unit size distribution

Primary-unit radii were sampled from a lognormal distribution with a prescribed coefficient of variation. Following sampling, all radii in each realization were rescaled such that

$$\left\langle a^3 \right\rangle^{1/3} = a_{\text{ref}}, \tag{5}$$

where $a_{\mathrm{ref}}$ is the reference primary-unit radius. Consequently, the average primary-unit volume, and hence the average primary-unit mass, remains constant between realizations. This prevents changes in aggregate scaling from arising trivially from changes in the average primary-unit mass.

For the simulations used to calibrate the relationship between fusion extent and fractal dimension, primary-unit radii were sampled with a coefficient of variation CV = 0.5.

###### 1.6.2 Brownian cluster–cluster aggregation

Each simulation was initialized with $N$ isolated primary units. Aggregation proceeded irreversibly by repeatedly selecting two existing clusters, bringing them into first contact, and replacing them with the resulting merged cluster.

Cluster pairs were selected using a Smoluchowski-like Brownian collision kernel,

$$K_{ij} \propto (D_i + D_j)\,(R_i + R_j)\,, \tag{6}$$

where $R_i$ is the collision scale of cluster $i$ and its translational diffusion coefficient was approximated as $D_i \propto R_i^{-1}$. Pair-selection probabilities therefore account for both the size-dependent Brownian mobility and the effective collision cross-section of the two clusters.

The geometry of individual encounters was determined using Brownian first-passage capture. One cluster was launched outside the other and propagated by translational Brownian motion until the first contact between their constituent spherical elements. Rotational Brownian diffusion of the incoming cluster was included during the encounter. The production simulations used a bounding-volume-hierarchy accelerated implementation of the Brownian first-passage procedure, including a heuristic rotational diffusion scale. This capture procedure preferentially samples exposed regions of open aggregates, thereby incorporating the screening characteristic of diffusion-limited aggregation.

###### 1.6.3 Geometric representation of incomplete fusion

Following first contact, the degree of fusion was controlled by a dimensionless fusion extent $f$, with

$$0 \le f \le 1. \tag{7}$$

At $f = 0$, the contacting spherical elements remain at their hard-sphere touching distance. For finite $f$, the separation between the contacting subclusters is reduced according to

$$d_{\mathrm{target}} = (1 - f)d_{\mathrm{touch}} + f d_{\mathrm{coal}}, \tag{8}$$

where $d_{\mathrm{touch}}$ is the center-to-center distance at first contact and $d_{\mathrm{coal}}$ is the separation at which the two spherical mass elements have the same combined radius of gyration as a single volume-equivalent sphere. The two colliding subclusters are translated toward one another while preserving their combined center of mass.

After formation of the final aggregate, intermediate fusion is additionally represented by an isotropic relaxation of the spherical mass-element coordinates toward the radius of gyration of the corresponding volume-equivalent compact sphere. Total mass is conserved and a lower bound is imposed such that the aggregate cannot become more compact than a homogeneous sphere containing the same total volume. At $f = 1$, the limiting structure is a single volume-equivalent sphere.

The fusion extent $f$ should therefore be regarded as a geometric measure of relaxation toward complete coalescence, rather than representing any physical quantity. The model tests whether varying the degree of incomplete fusion is sufficient to reproduce the experimentally observed mesoscale structure; it is not intended as a dynamical model of condensate coalescence.

###### 1.6.4 Radius of gyration and mass–size scaling

For an aggregate containing spherical elements with masses $m_i \propto a_i^3$ and mass-centered positions $\mathbf{r}_i$, the radius of gyration was calculated as

$$R_g^2 = \frac{\sum_i m_i \left(|\mathbf{r}_i|^2 + \frac{3}{5}a_i^2\right)}{\sum_i m_i}. \tag{9}$$

The term $3a_i^2/5$ accounts for the finite internal radius of gyration of each spherical primary unit. Aggregate mass was calculated additively as

$$M \propto \sum_i a_i^3. \tag{10}$$

The simulated structures were characterized through the mass–size relation

$$\frac{M}{m_{\text{ref}}} = k_f \left(\frac{R_g}{a_{\text{ref}}}\right)^{D_f}, \tag{11}$$

where $m_{\text{ref}} \propto a_{\text{ref}}^3$, $D_f$ is the effective fractal dimension, and $k_f$ is the fractal prefactor. For each number of primary units $N$, independent aggregate realizations were generated and the median values of $M$ and $R_g$ were used for fitting. The fractal dimension was obtained as the slope of a linear regression of

$$\ln\left(\frac{M}{m_{\text{ref}}}\right) = \ln k_f + D_f \ln\left(\frac{R_g}{a_{\text{ref}}}\right). \tag{12}$$

Bootstrap confidence intervals were obtained by resampling independent aggregate realizations within each value of $N$, recalculating the size medians, and repeating the scaling fit.

For the calibration used to establish the relationship between $D_f$ and fusion extent, simulations were performed at

$$f = 0,\ 0.4,\ 0.7,\ \text{and } 0.9, \tag{13}$$

using primary-unit CV $= 0.5$. Aggregate sizes of

$$N = 16,\ 32,\ 64,\ \text{and } 128 \tag{14}$$

primary units were simulated, with 32 independent realizations for each aggregate size and fusion extent. Increasing $f$ generated progressively more compact aggregates and a corresponding increase in the fitted effective fractal dimension. This relationship was used in Fig. 4c to compare the experimentally determined fractal dimensions with the degree of relaxation in the partial-fusion model.

##### 1.6.5 Simulated scattering profiles

To determine whether the simulated aggregate structures can also reproduce the characteristic scattering signatures of the fuzzy assemblies, rotationally averaged scattering profiles were calculated within the Rayleigh–Debye–Gans approximation.

The amplitude form factor of a homogeneous spherical mass element of radius $a$ is

$$F(qa) = 3\frac{\sin(qa) - qa\cos(qa)}{(qa)^3}. \tag{15}$$

For an aggregate, the orientationally averaged scattering intensity was calculated from the coherent Debye sum,

$$I(q) = \sum_{i,j} b_i(q) b_j^*(q) \frac{\sin(qr_{ij})}{qr_{ij}}, \tag{16}$$

where $r_{ij}$ is the distance between scattering elements $i$ and $j$. Assuming a common optical contrast,

$$b_i(q) \propto a_i^3 F(qa_i). \tag{17}$$

The calculated scattering curves were normalized by the forward intensity $I(0)$. For comparison with the experimentally measured scattering amplitude, the square root of the normalized intensity was used,

$$\frac{|A(q)|}{|A(0)|} = \sqrt{\frac{I(q)}{I(0)}}. \tag{18}$$

The Rayleigh–Debye–Gans calculation neglects multiple scattering, near-field coupling, and shadowing between the spherical elements. It is therefore used here to assess qualitative form-factor features arising from the simulated mass distribution rather than to predict absolute scattering intensities. In particular, partially fused non-compact aggregates display suppression of the pronounced spherical form-factor minimum, qualitatively reproducing the characteristic scattering behavior of the experimentally identified fuzzy assemblies.

##### 1.6.6 Hydrodynamic properties of simulated aggregates

For comparison with the simultaneously measured optical and hydrodynamic radii, the translational hydrodynamic radius of each simulated aggregate was calculated from the rigid-body mobility of its constituent spherical elements using a Rotne–Prager–Yamakawa mobility representation. The resulting $R_h$ corresponds to the radius of an equivalent sphere with the same translational hydrodynamic mobility.

For the hydrodynamic-size analysis, additional simulations were performed at primary-unit CV = 0.5 for fusion extents $f = 0$ and $f = 0.9$. Aggregate sizes of

$$N = 8,\ 16,\ 32,\ 64,\ 128,\ 256,\ \text{and } 512 \tag{19}$$

were simulated with 16 independent realizations per aggregate size and fusion extent. Eight additional aggregates containing $N = 1024$ primary units were generated for $f = 0.9$. These simulations used the same Brownian pair-selection and Brownian first-passage capture procedure as the fractal-dimension simulations, with rotational diffusion enabled.

#### 2 Supporting figures

The supporting figures provide additional datasets and control measurements that complement the main-text analysis. These data demonstrate the robustness and reproducibility of the observed trends in particle size, refractive index, and interfacial structure across independent condensate preparations and across distinct polymer systems.

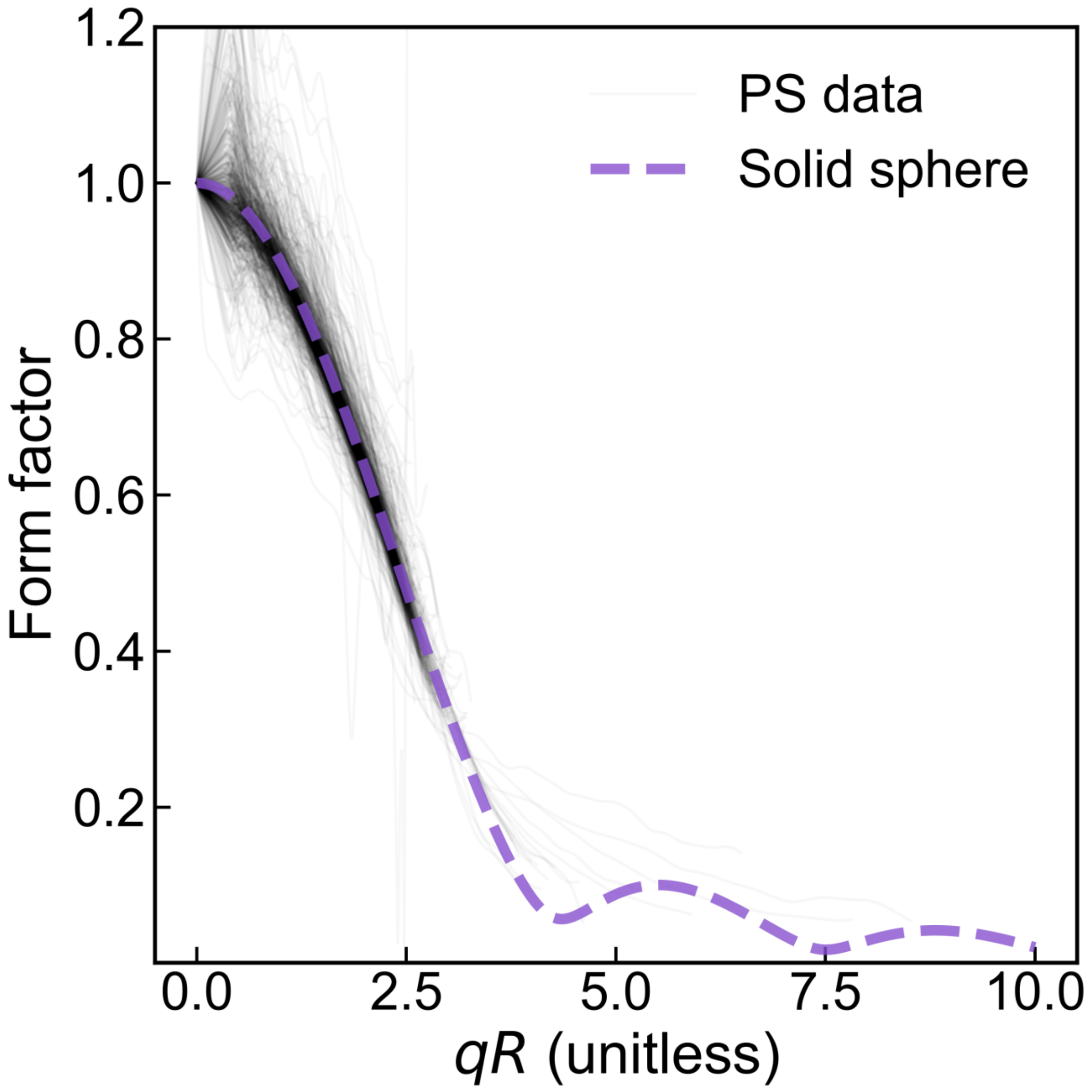


Figure 1: **Validation of form-factor analysis using polystyrene (PS) reference particles.** Experimental scattering form factor as a function of $qR$, where $q$ is the magnitude of the scattering wave vector and $R$ is the particle radius. Solid black lines represent experimental data for polystyrene particles, and the dashed purple curve shows the theoretical Mie scattering prediction.

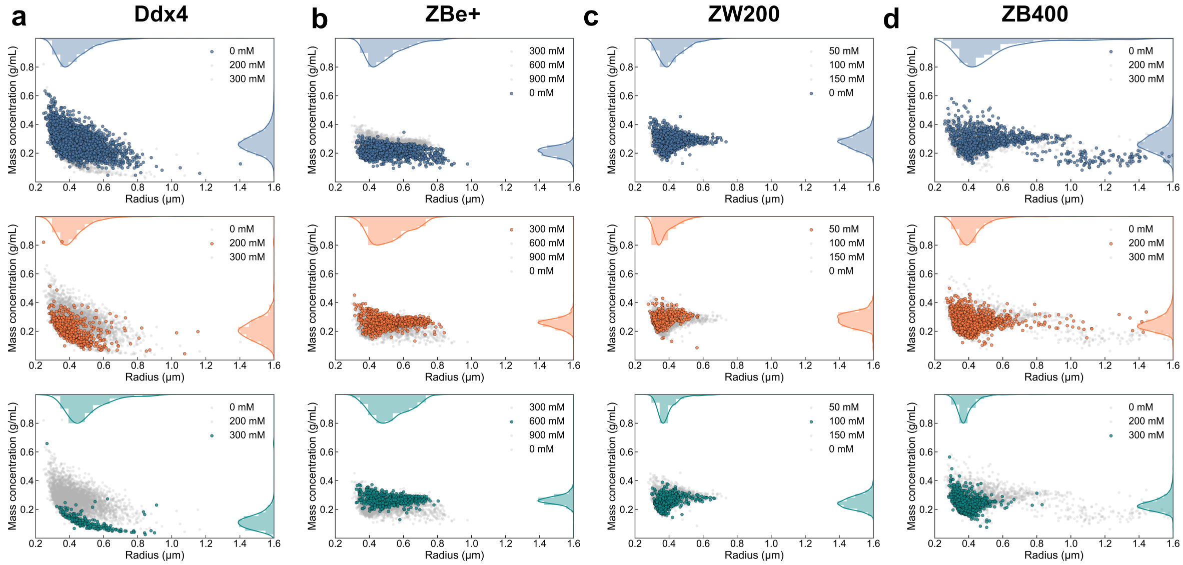


Figure 2: **Salt dependence of inferred mass concentration for additional condensate and polymer systems.** (**a**) Ddx4 condensates, (**b**) ZBe+ polymer condensates, (**c**) ZW200 polymer condensates, and (**d**) ZB400 polymer condensates. Each panel shows the inferred mass concentration as a function of optical radius for thousands of individual condensates measured at multiple salt concentrations, as indicated in the legends. Ddx4 condensates exhibit a pronounced size dependence in mass concentration and a clear decrease in concentration with increasing ionic strength. In contrast, the polymer condensates, particularly ZBe+ and ZW200, display a weaker size dependence and reduced sensitivity to salt, consistent with more uniform internal packing across the conditions investigated.

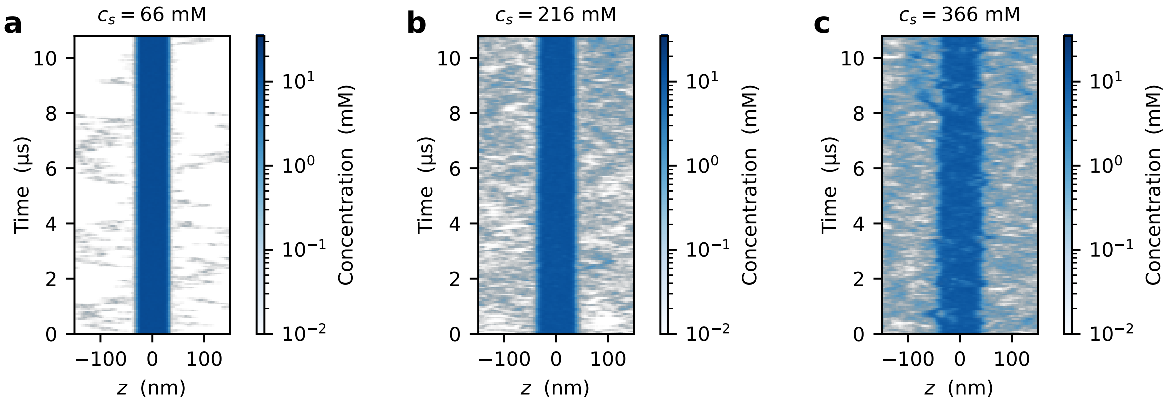


Figure 3: **Time series of the protein concentration from molecular dynamics simulations:** Ddx4N1 concentration profiles along the long axis of the simulation box at salt concentration, $c_s$, (**a**) 66 mM, (**b**) 216 mM, and (**c**) 366 mM.

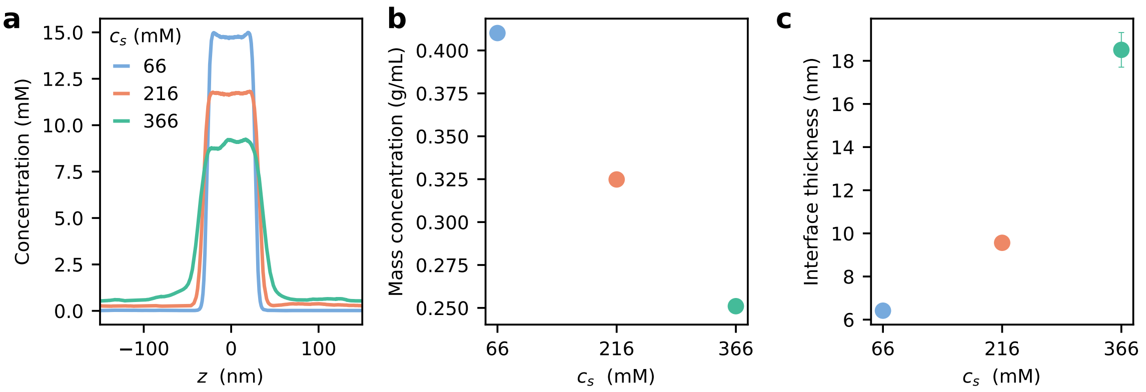


Figure 4: **Interface thickness increases with salt concentration, $c_s$, in molecular dynamics simulations:** (**a**) Protein concentration along the long axis of the simulation box. (**b**) Comparison of mass concentrations of the protein-rich phase from simulations (circles) and experiments (squares). (**c**) Interfacial widths estimated from simulations at different $c_s$ values.

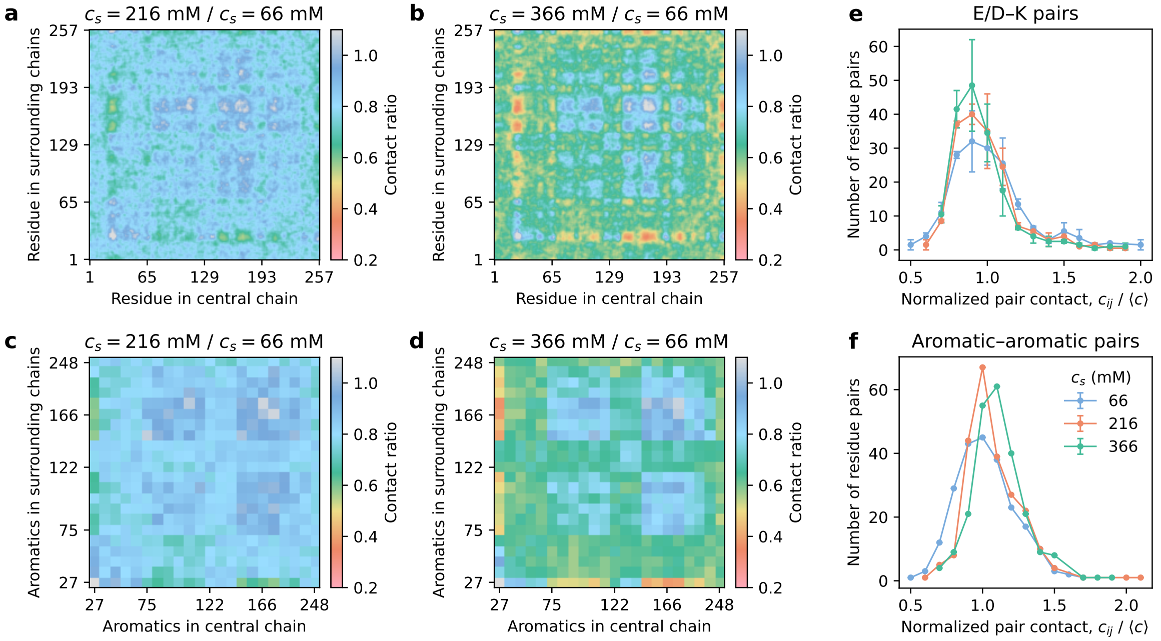


Figure 5: **Intermolecular contact maps show a relative increase in interactions between aromatic residues with increasing salt concentrations, $c_s$:** (**a**–**b**) Contact maps for all residues calculated for a protein chain in the center of the condensate and the surrounding chains at $c_s$ (**a**) 216 mM and (**b**) 366 mM. (**c**–**d**) Contact maps for only aromatic residues at $c_s$ (**c**) 216 mM and (**d**) 366 mM. Contact maps in **a**–**d** are normalized by the contact map at $c_s$ = 66 mM. (**e**–**f**) Distributions of contacts between (**e**) oppositely charged residues and (**f**) aromatic residues at different $c_s$ values. To account for the decrease in dense-phase concentrations with increasing $c_s$, contacts in **e**–**f** are normalized by the average number of contacts between all residues at the corresponding $c_s$.

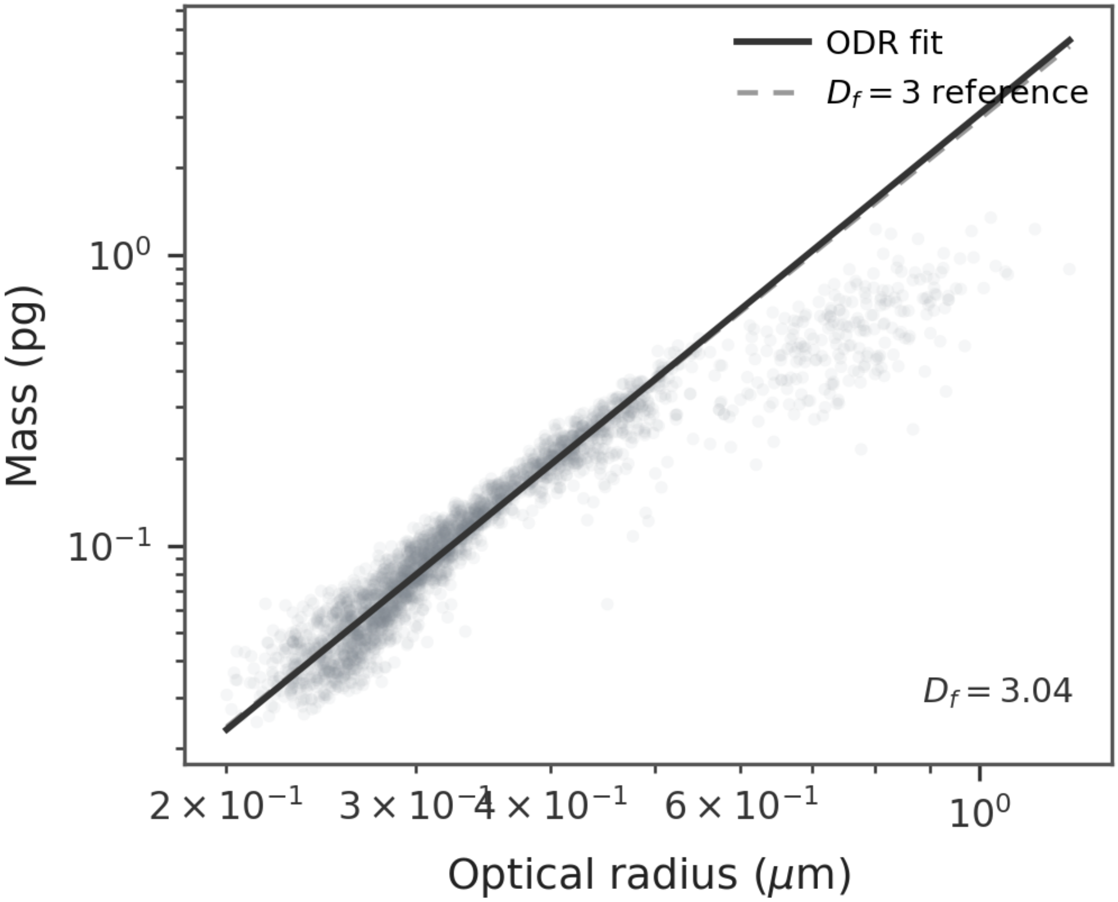


Figure 6: **Mass-size scaling of sharp assemblies** Sharp condensates ($\rho^2/R^2 < 0.2$) display scaling consistent with homogeneous spheres ($D_\mathrm{f} \sim 3$). The data corresponds to 66 mM NaCl, as this was the only condition in which the sharp population spanned a sufficiently large size range to allow accurate fitting of scaling exponent.

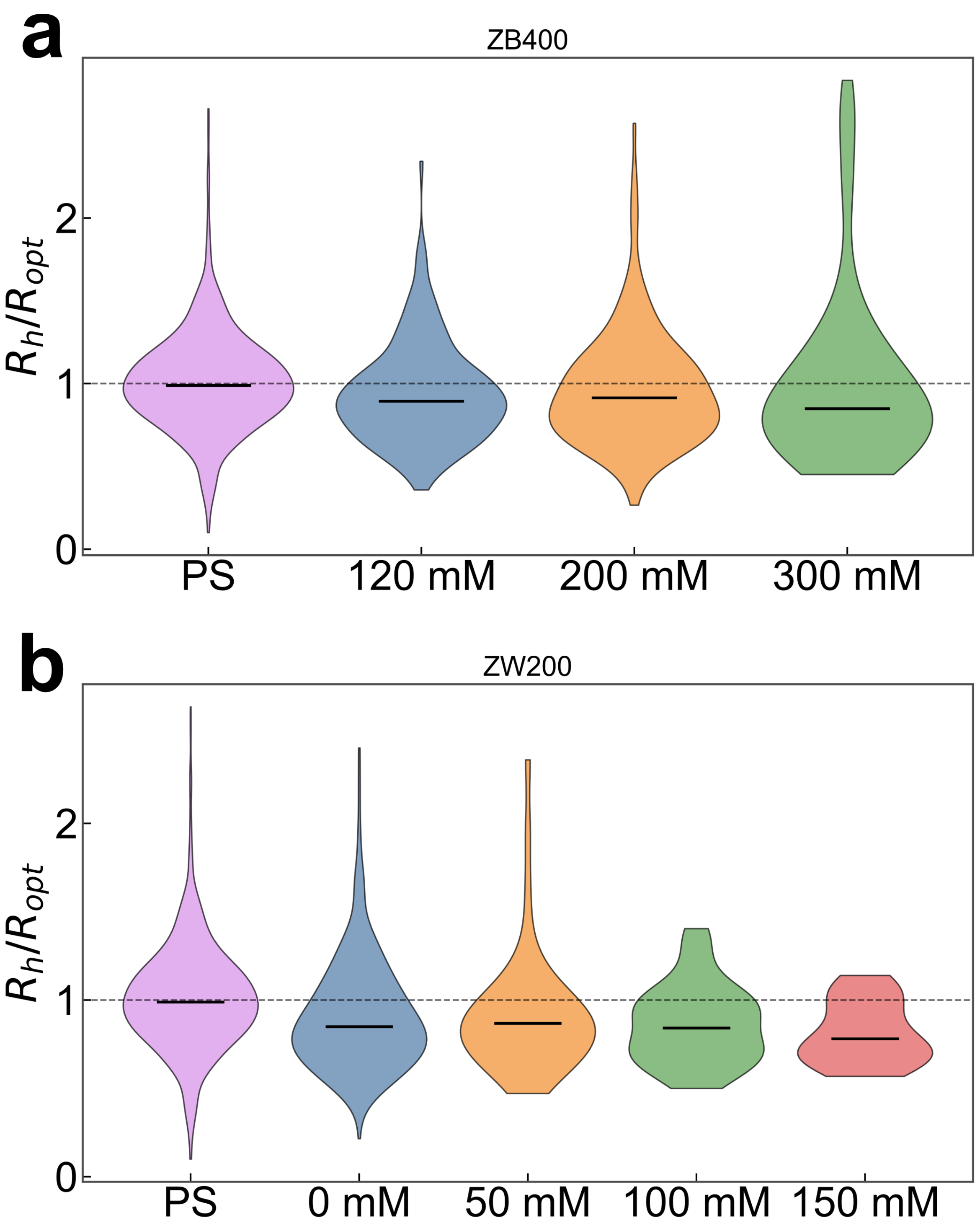


Figure 7: **Distribution of the ratio of hydrodynamic radius to optical radius.** (**a**)ZB condensates measured at different salt concentrations (120, 200, and 300 mM). (**a**) Distribution of the ratio of hydrodynamic radius to optical radius ($R_\mathrm{h}/R_\mathrm{opt}$) for individual condensates. Polystyrene (PS) particles are shown as a reference. The dashed line indicates $R_\mathrm{h} = R_\mathrm{opt}$. (**b**) Equivalent analysis for ZW200 condensates measured at 0, 50, 100, and 150 mM salt. **b** Distribution of $R_\mathrm{h}/R_\mathrm{opt}$ for individual condensates compared with PS reference particles.
